# Chromosome-scale assembly and annotation of the macadamia genome (*Macadamia integrifolia* HAES 741)

**DOI:** 10.1101/2020.05.25.114009

**Authors:** Catherine J. Nock, Abdul Baten, Ramil Mauleon, Kirsty S. Langdon, Bruce Topp, Craig Hardner, Agnelo Furtado, Robert J. Henry, Graham J. King

## Abstract

*Macadamia integrifolia* is a representative of the large basal eudicot family Proteaceae and the main progenitor species of the Australian native nut crop macadamia. Since its commercialisation in Hawaii fewer than 100 years ago, global production has expanded rapidly. However, genomic resources are limited in comparison to other horticultural crops. The first draft assembly of *M. integrifolia* had good coverage of the functional gene space but its high fragmentation has restricted its use in comparative genomics and association studies. Here we have generated an improved assembly of cultivar HAES 741 (4,094 scaffolds, 745 Mb, N50 413 kb) using a combination of Illumina paired and PacBio long read sequences. Scaffolds were anchored to 14 pseudo-chromosomes using seven genetic linkage maps. This assembly has improved contiguity and coverage, with >120 Gb of additional sequence. Following annotation, 34,274 protein-coding genes were predicted, representing 92% of the expected gene content. Our results indicate that the macadamia genome is repetitive and heterozygous. The total repeat content was 55% and genome-wide heterozygosity, estimated by read mapping, was 0.98% or one SNP per 102 bp. This is the first chromosome-scale genome assembly for macadamia and the Proteaceae. It is expected to be a valuable resource for breeding, gene discovery, conservation and evolutionary genomics.

## Introduction

The genomes of most crop species have now been sequenced and their availability is transforming breeding and agricultural productivity^1,2^. Macadamia is the first Australian native plant to become a global food crop. In 2018/2019, world production was valued at US$1.1 billion (59,307 MT/year, kernel basis), reflecting the most rapid increase in production of any nut crop over the past 10 years^3^. As a member of the Gondwanan family Proteaceae (83 genera, 1660 species)^4^, *Macadamia* (F.Muell.) is a basal eudicot and is phylogenetically divergent from other tree crops^5^. *M. integrifolia*, the main species used in cultivation^6^, is a mid-storey tree endemic to the lowland rainforests of subtropical Australia^7^. Macadamia was commercialised as a nut crop from the 1920s in Hawaii and cultivated varieties are closely related to their wild ancestors in Australia^8^.

Despite the rapid expansion in production over the past 50 years and commercial cultivation in 18 countries, breeding is restricted by a paucity of information on the genes underlying important crop traits^9^. There are few genomic resources for either macadamia or within the Proteaceae. Transcriptomic data are available for *Macadamia*^10^ and other Proteaceae genera including *Banksia*^11^, *Grevillea*^12^, *Protea*^13^ and *Gevuina*^14^, and have been utilised to help to understand the evolution of floral architecture^12^ and variation in locally adaptive traits^13^. Recent genome wide association studies in macadamia have identified markers associated with commercially important traits^15^ but their location and context in the genome is unknown. An earlier highly fragmented (193,493 scaffolds, N50 4,745)^16^ draft genome assembly of the widely grown *M. integrifolia* cultivar HAES 741 was constructed from Illumina short read sequence data.

Here we report on the first annotated and anchored genome assembly for macadamia and the Proteaceae. Long read PacBio and paired-end Illumina sequence data were generated and 187 Gb of data were used to construct an improved HAES 741 assembly (Table 1). Assembled sequence scaffolds from the new hybrid *de novo* assembly (745 Mb, 4094 scaffolds, N50 413 kb) were then anchored and oriented to 14 pseudo-chromosomes using methods to maximise colinearity across seven genetic linkage maps^17^. A contiguous genome sequence for macadamia is expected to facilitate the identification of candidate genes and support marker assisted selection to accelerate the development of new improved varieties^9^.

**Table 1:** Data files and library information for *Macadamia integrifolia* genome sequencing. *Data deposited for draft assembly v1.1

| Sequencing Platform | Library | Insert size (bp) | Read length (bp) | Sequence reads (Million) | Sequence bases (Gb) | Accession |
| --- | --- | --- | --- | --- | --- | --- |
| Genomic data |  |  |  |  |  |  |
| Illumina | Pair end | 200 | 2 x 125 | 228.8 | 57.2 | SRR10896963 |
|  | Pair end | 350 | 2 x 125 | 119.0 | 29.8 | SRR10896962 |
|  | Pair end | 550 | 2 x 125 | 107.6 | 26.9 | SRR10896961 |
|  | Pair end | 480 | 2 x 150 | 58.7 | 17.7 | ERX1468522-23* |
|  | Pair end | 700 | 2 x 150 | 28.7 | 8.7 | ERX1468524* |
|  | Mate pair | 8000 | 2 x 100 | 200.3 | 40.5 | ERX1468525* |
| PacBio | NA |  | 9769 | 1.0 | 6.4 | SRR10896960 |
| Total gDNA |  |  |  | 744.1 | 187.2 |  |
| Transcriptomic data |  |  |  |  |  |  |
| Illumina | young leaf | 300 | 2 x 100 | 77.9 | 15.6 | SRR10897159 |
|  | shoot | 300 | 2 x 100 | 71.6 | 14.3 | SRR10897158 |
|  | flower | 300 | 2 x 100 | 83.3 | 16.7 | SRR10897157 |
| Total RNA-seq |  |  |  | 232.8 | 46.6 |  |

## Materials and Methods

### Sample collection and extraction

Leaf, shoot and flower tissue were collected from a single *M. integrifolia* cultivar HAES 741 individual located in the M2 Regional Variety Trial plot, Clunes, New South Wales, Australia (28°43.843’S; 153°23.702’E). An herbarium specimen was submitted to the Southern Cross Plant Science Herbarium [accession PHARM-13-0813]. On collection, samples were snap frozen in liquid nitrogen or placed on dry ice and stored at −80ºC prior to extraction. Previously described methods for and DNA/RNA extraction for Illumina^16^ and PacBio^18^ sequencing were followed.

### Library preparation and sequencing

In addition to the original 480 bp and 700 bp insert and 8 kb mate pair libraries (51.6 Gb total), new libraries with 200, 350 and 550 bp insert sizes were prepared with TruSeq DNA PCR-free kits and sequenced with Illumina HiSeq 2500 producing 57.0, 29.8 and 29.6 Gb paired-end sequence data. A PacBio library (20-Kb) was prepared and sequenced across 10 SMRT cells on a PacBio RSII system (P6-C4 chemistry) generating 6.44 Gb data. Transcriptome sequencing of leaf, shoot and flower tissue followed previously described methods^16^ and generated 44.6 Gb paired-end RNA-seq data in total (Table 1).

### De novo assembly and anchoring

Illumina raw read data were initially assessed for quality using FastQC^19^. Low quality bases (Q<20), adapters and chloroplast reads were removed from Illumina data using BBMaps^20^. PacBio reads were error-corrected using the LoRDEC^21^ hybrid error correction method. Hybrid *de novo* assembly, scaffolding and gap closing was performed using MaSuRCA^22^. Illumina paired-end and mate-pair reads were extended into super-reads of variable lengths, and combined with PacBio reads to generate the assembly. The MaSURCA output was further scaffolded in two rounds, firstly using SSPace^23^ scaffolder with the Illumina mate pair reads, and secondly, using L_RNA_Scaffolder^24^ with the long transcripts generated using a Trinity^25^ pipeline and RNA-Seq reads. Scaffolds were anchored and oriented using ALLMAPS^26^ with 4,266 ordered sequence-based markers across seven genetic linkage maps generated from four mapping populations with HAES 741 parentage^17^.

### Genome size and heterozygosity

For genome size and heterozygosity estimation, k-mer frequency was determined from cleaned short and long read data for k ranging from 15 to 33 at a step of 2 in Jellyfish^27^. Genome size was estimated at the optimal k-mer of 25, with short read data only using findGSE because it is an optimal method for plants^28^. Genome-wide heterozygosity using reads data was estimated with GenomeScope^29^ from the k-mer 25 histogram computed using Jellyfish. In addition, genome-based heterozygosity was determined by mapping post-QC short read data to the assembly using minimap2^30^ and heterozygous sites identified using GATK best practices^31^ for SNP discovery.

### Genome annotation and gene prediction

Repetitive elements were first identified in the final assembly by modelling repeats using RepeatModeler, and then quantified using RepeatMasker^32^. Transcriptome assembly was performed using the Trinity pipeline^25^. Following repeat masking, the final assembly was annotated using the MAKER gene model prediction pipeline^33^. Sources of evidence for gene prediction included the Trinity assembled transcripts and protein sequences of the taxonomically closest available genome sequence of the sacred lotus *Nelumbo nucifera*, and the model plant *Arabidopsis thaliana*.

### Comparative analysis of orthologous eudicot genes

Orthologous gene clusters were identified and compared using OrthoVenn2^34^ and protein sequences from *M. integrifolia*, *N. nucifera*, *A. thaliana*, *Prunus persica* (peach), *Eucalpytus grandis* and *Coffea canephora* (coffee) genomes. Pairwise sequence similarities were determined applying a BLASTP E-value cut-off of 1E-05 and an inflation value of 1.5 for OrthoMCL Markov clustering. Macadamia specific gene clusters were tested for GO enrichment using OrthoVenn2.

### Quality Assessment

The completeness of the genome assembly was evaluated by BUSCO (benchmarking universal single copy orthologs)^35^ using the green plant dataset (viridiplantae.odb10). To further assess the accuracy of the *M. integrifolia* genome assembly, short reads were mapped to the pseudo-genome using Minimap2^30^ and the number of mapped reads were computed using paftools.js.

### Data Availability

The *M. integrifolia* Whole Genome Shotgun project has been deposited at DDBJ/ENA/GenBank under the accession JAAEEG000000000. The version described in this paper is JAAEEG010000000. Datasets generated in this study have been deposited at NCBI under BioProject number PRJNA593881^36^. Raw genomic DNA and RNA-seq read files have been deposited in the NCBI Sequence Read Archive (Table 1).

## Results and Discussion

### Genome Assembly

Hybrid *de novo* assembly using 180.8 Gb Illumina short read and 6.44 Gb PacBio long read data produced a 744.6 Mb genome assembly (v2) with 180 x coverage of the genome and a scaffold N50 of 413 kb (Table 1). In comparison to the v1.1 assembly of the same cultivar, this v2 assembly represents an ~68 fold increase in contiguity and includes 120.3 Mb of new sequence (Table 2). Scaffolds were anchored and oriented to 14 pseudo-chromosomes using ALLMAPs to maximise the collinearity of ordered sequence-based markers across seven genetic linkage maps generated from mapping populations with HAES 741 parentage^17^. This anchored 69.7% of the assembly to 14 pseudo-chromosomal sequences ranging in size from 29.2 to 47.0 Mb (Table 3). The quality and completeness of the assembly, assessed using BUSCO, indicates that the macadamia assembly contains 91.9% of the expected single copy gene content. In addition, the accuracy of the assembly was assessed by short read mapping with 99% of reads (98.97-99.15% per library) aligning reliably to the assembly.

**Table 2:** Comparison of the new *Macadamia integrifolia* cv. HAES 741 genome assembly with the previously published draft assembly. Coverage is based on k-mer estimated genome size of 895.7 Mb

|  | Data |  | Scaffold |  |  | Assembly |  |  | Annotation |  |  |
| --- | --- | --- | --- | --- | --- | --- | --- | --- | --- | --- | --- |
|  | Post-QC Gb | Genome Coverage X | Number | N50 kb | Longest Mb | Assembled Genome Mb | Gaps Mb | Genome Coverage % | Repeat Content | Gene Models | BUSCO |
| v1.1 | 51.6 | 58 | 193,493 | 4.7 | 0.64 | 518.49 | 70 | 58% | 37% | 35,337 | 77.4% |
| **v2** | 161.5 | **180** | **4,094** | 413 | 2.19 | 744.64 | 11 | **83%** | 55% | 34,274 | **91.9%** |

**Table 3:** Summary of the assembled chromosomes of macadamia

| Chromosome | Length<br>bp | scaffolds | genes |
| --- | --- | --- | --- |
| Chr01 | 36,399,236 | 126 | 1543 |
| Chr02 | 44,010,915 | 120 | 2214 |
| Chr03 | 37,831,390 | 112 | 2039 |
| Chr04 | 37,507,012 | 91 | 2133 |
| Chr05 | 46,991,412 | 110 | 2334 |
| Chr06 | 40,842,004 | 101 | 1941 |
| Chr07 | 36,789,931 | 112 | 1940 |
| Chr08 | 34,869,527 | 94 | 1975 |
| Chr09 | 42,247,492 | 109 | 2019 |
| Chr10 | 34,167,173 | 87 | 1701 |
| Chr11 | 33,967,801 | 107 | 1488 |
| Chr12 | 31,654,987 | 87 | 1523 |
| Chr13 | 29,219,948 | 100 | 1398 |
| Chr14 | 32,845,142 | 109 | 1531 |
| **Anchored** | **519,343,970** | **1465** | **25779** |
| Unplaced | 225,433,603 | 2629 | 8,495 |
| **Total** | **744,777,573** | **4094** | **34,274** |

### Genome size, heterozygosity and repetitive content

Genome size estimation of 896 Mb was 1.37 times larger than the only previous estimate for *M. integrifolia* of 652 Mb (600-700 Mb) that was based on 51.6 Gb Illumina short read data^(16)^. The new k-mer based estimate is considered to be more accurate due to improved read coverage from an additional 114 Gb of high-quality Illumina data. Based on the revised genome size estimate, the v2 assembly covers 83% of the genome. Average genome heterozygosity, determined using short read data from the k-mer 25 Jellyfish^(27)^ histogram, was 1.36% (Fig. 2). Genome-wide heterozygosity was slightly lower at 0.98% for the assembly-based SNP analysis method^30, 31^. In total, 7,309,539 heterozygous sites were identified in the HAES 741 genome assembly representing one SNP per 102 bp. This is consistent with reports that macadamia is highly heterozygous and predominantly outcrossing^9^. Repetitive content accounted for 410.5Mb (55.1%) of the assembly. As reported for many other plant genomes, long terminal repeat (LTR) retrotransposons were the most abundant repeat type (Fig. 3).

**Figure 1.**
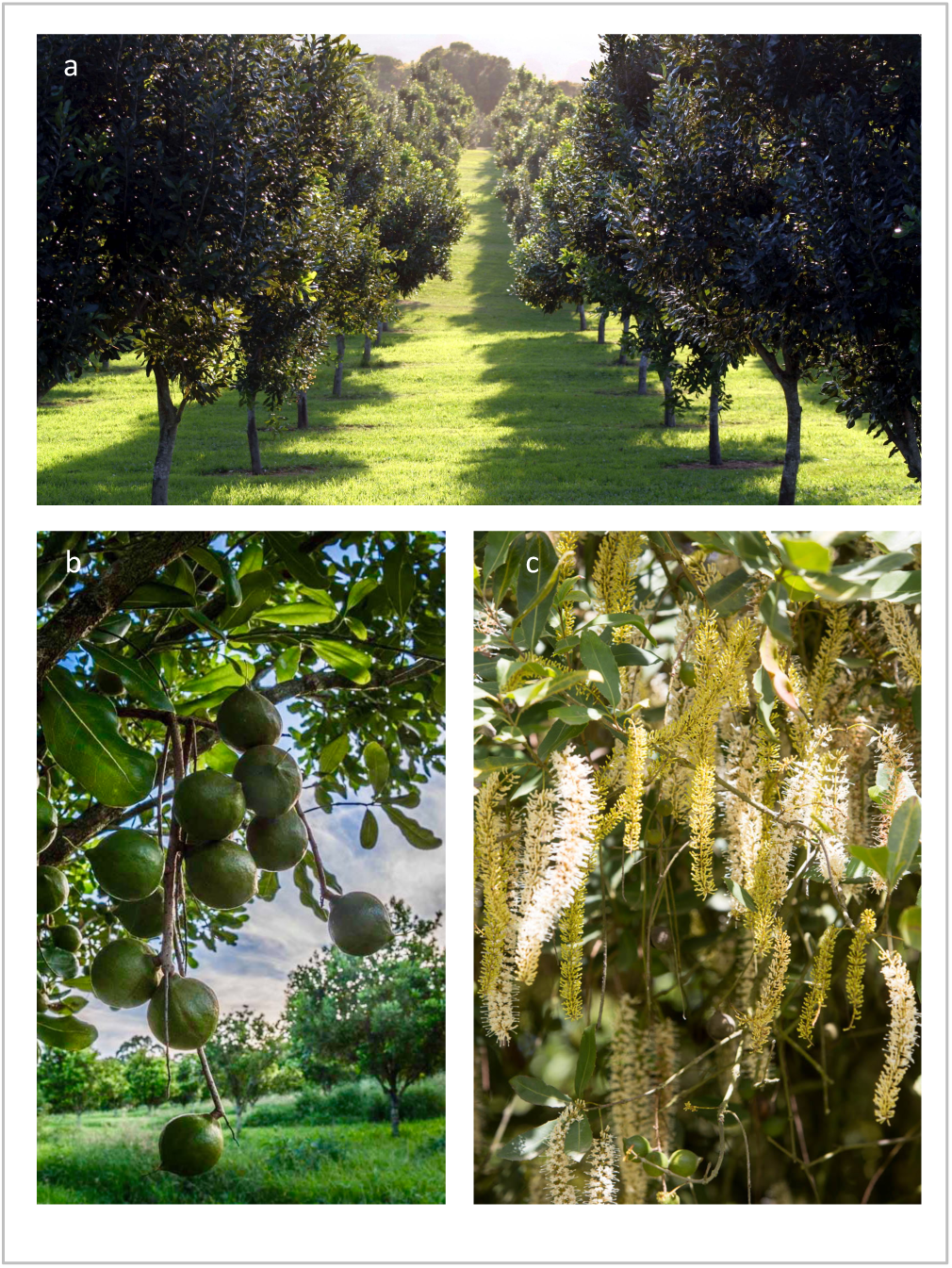
*Macadamia integrifolia* (a) orchard (b) nut in husk (c) racemes

**Figure 2.**
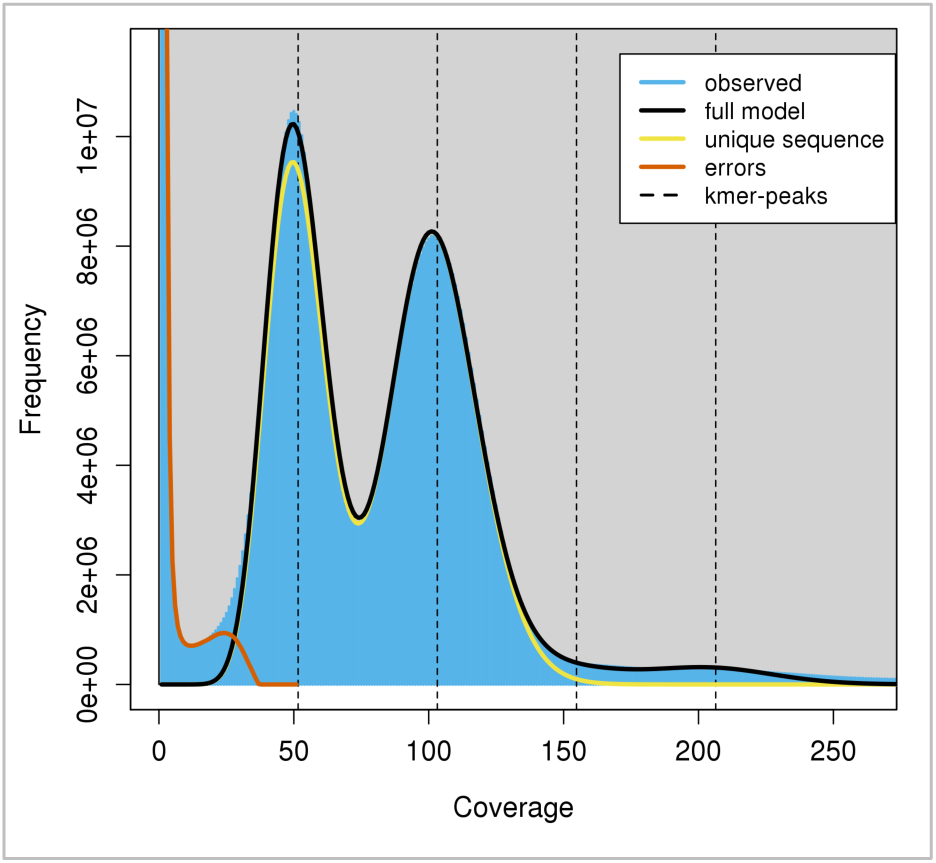
The 25-mer distribution for estimation of genome heterozygosity and size. Peaks at approximately 50, 100 and 200 represent heterozygous, homozygous and repeated k-mers respectively.

**Figure 3.**
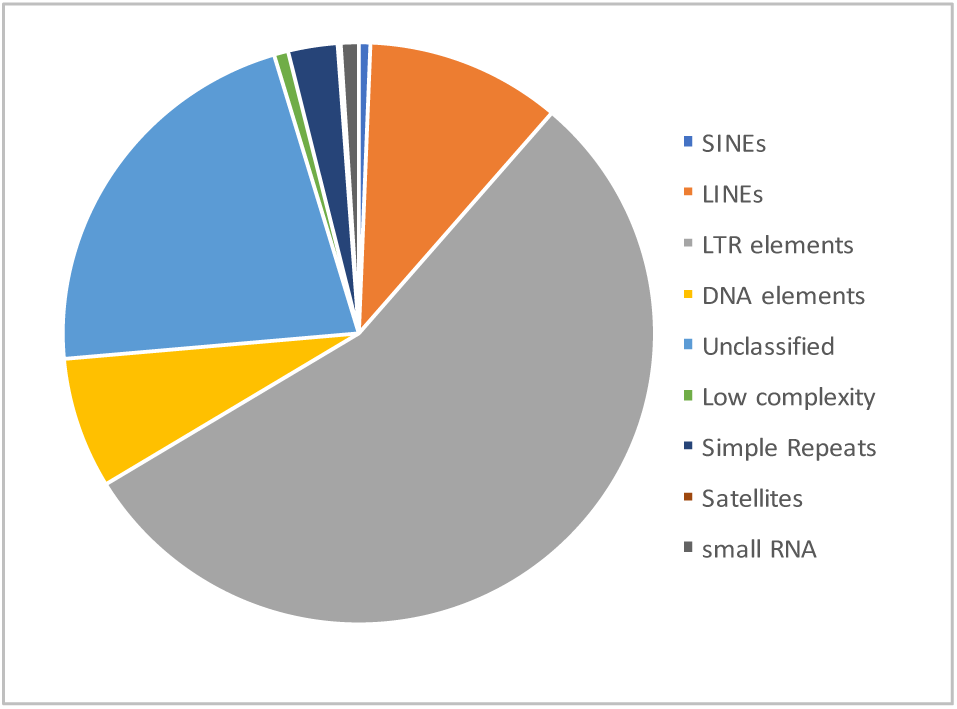
Repetitive elements representing 55.1% of the macadamia genome assembly

### Annotation and comparative analysis of orthologous genes

Annotation of the final assembly, identified 34,274 high-confidence gene models. Of these, 25,779 (75.2%) were anchored to pseudo-chromosomes and 8,495 were located on unanchored scaffolds (Table 3). Comparative analysis of the macadamia gene models with the proteins of *N. nucifera*, *A. thaliana*, *P. persica*, *E. grandis* and *C. canephora* identified 8,961 orthologous clusters including 2,051 single gene clusters that were shared by all six eudicot species (Fig 4). *M. integrifolia* and *N. nucifera* both belong to the basal eudicot order Proteales and contained similar numbers of clusters with 13,491 and 13,321 respectively. Tests for gene ontology (GO) enrichment of 1,094 macadamia specific clusters identified five significant terms including those associated with the defence response (GO:0006952, P = 4.8E-05) and fruit ripening (GO:0009835, P = 5.9E-04).

**Figure 4.**
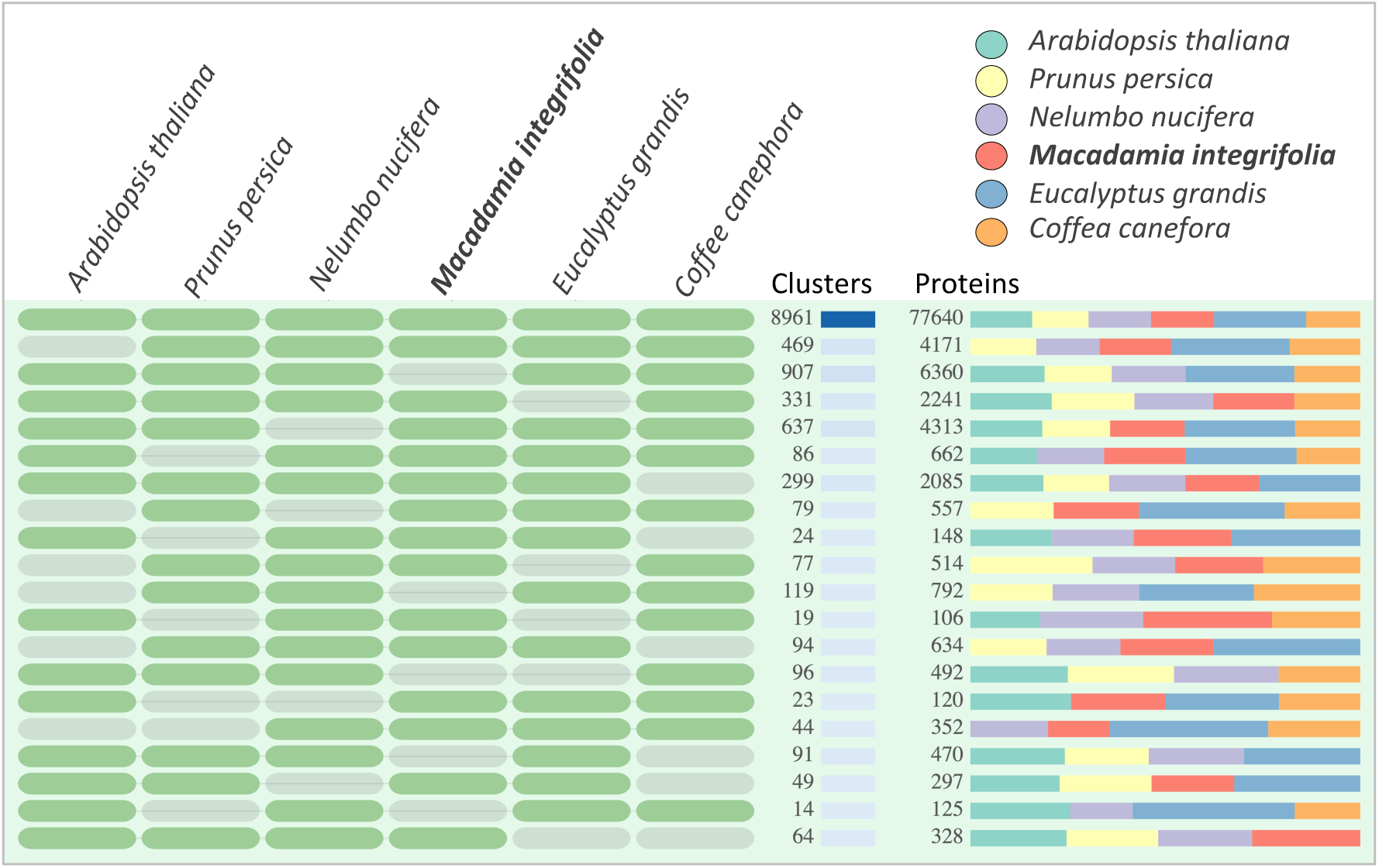
Comparison of orthologous gene families (clusters) and proteins among six eudicot species. On left, the groups of species included in each of 20 comparisons are shown in dark green with the corresponding number of orthologous clusters and proteins for each comparison. On right, the relative proportions of proteins for each species.

### Conclusion

Here we present the first chromosome-scale genome assembly for the nut crop macadamia and the large Gondwanan plant family Proteaceae. This provides a platform for unravelling the genetics of macadamia and is expected to underpin future breeding, and comparative and horticultural genomics research.

### Code Availability

Versions and parameters of the tools implemented in data analysis are provided below:

1. BBMap version 36.62: *ktrim=r k=23 mink=11 hdist=1 tpe tbo maxns=1 minlen=50 maq=8 qtrim=rl trimq=20*
2. LoRDEC version 0.4: *lordec-correct −2 input_for_read_correction.fastq -k 19 -S out.stat.txt -s 3 -T 12 -i PacBio_filtered_reads.fasta -o out.pacbio.corrected.fasta*
3. MaSuRCA version 3.2.1. Default parameters except: *NUM_THREADS = 16, JF_SIZE = 20000000000* (jellyfish hash size)
4. L_RNA_scaffolder: *blat -t=dna -q=dna scaffolds_gapClosed_min1000.fa Trinity.fasta transcript.vs.macaAssembly.psl -noHead -out=psl*
5. ALLMAPS version 0.7.7: Default parameters
6. Jellyfish version 2.0: *jellyfish count -t 14 -C -m 27 -s 8G -o 27mer_maca_illumOnly_out <all Illumina-only WGS fastqs>jellyfish histo -t14 27mer_maca_illumOnly_out> 27mer_maca_illumOnly_out.histo*
7. findGSE: Default parameters
8. GenomeScope version 2.2.6:*kmer length 27, read length 125bp, Max kmer coverage 1000*
9. Trinity, version 2.0.3: *Trinity --seqType fq --max_memory 100G –left* *reads_S_R1_clean.fq,reads_F2_R1_clean.fq,reads_YL_R1_clean.fq --right* *reads_S_R2_clean.fq,reads_F2_R2_clean.fq,reads_YL_R2_clean.fq --CPU 8*
10. RepeatModeler version 2.0.1. Default parameters
11. RepeatMasker version 4.0.9. Default parameters
12. MAKER, version 2.31.10. Default parameters except: *Gene prediction methods Augustus and SNAP (trained with previously generated macadamia gene models); AED score = 0.40; Minimum protein length: = 50 amino acids*
13. BUSCO version 3.0.2: *busco -i proteins.fasta -l viridiplantae_odb10 -m proteins -o output_name*
14. Minimap2 version 2.17. Default parameters (paftool.js bundled with Minimap2): *minimap2 -ax sr ref_assemblyfasta fastq1stReadPair fastq2ndReadPair > ref_readPairs_aln.paf; k8 paftools.js stat ref_readPairs_aln.paf > alignment_mapstats*
15. GATK HaplotypeCaller version 4.1.4.1: *gatk --java-options “-Xmx8g” HaplotypeCaller -R refGenome.fasta-I input.bam -O output.g.vcf.gz -ERC GVCF -heterozygosity 0.01*
16. GATK GenotypeGVCFs version 4.1.4.1: *gatk --java-options “-Xmx4g” GenotypeGVCFs -R refGenome.fasta-V input.g.vcf.gz -O output.vcf.gz --heterozygosity 0.01*
17. GATK VariantsToTable version 4.1.4.1: *gatk-4.1.4.1/gatk --java-options “-Xmx8g” VariantsToTable -V output.vcf.gz -F CHROM -F POS -F TYPE -F REF -F ALT -F HET -GF GT -GF AD -O output_genoGVCF.table.txt*

## Acknowledgements

This research was funded by Horticulture Innovation Australia project MC15008 *Establishing an open-source platform for unravelling the genetics of Macadamia: integration of linkage and genome maps*. The project provided a PhD scholarship for K.L. with funding from Horticulture Innovation Australia, Knappick Foundation Ltd., Macadamia Conservation Trust, Australian Macadamia Society and Southern Cross University. Laboratory and horticultural support were provided by Asuka Kawamata, Tiffeny Byrnes and Alicia Hidden. Assistance with sample collection was provided by Dr Alam Mobashwer. We thank Kim Wilson and Alex Yong for providing access to the M2 regional variety trial site in Clunes, NSW, Australia where the HAES 741 tree used for genome sequencing is located.

## Authors’ contributions

C.N., G.K., B.T., C.H. and R.H. conceived and designed the study. C.N., A.B. and G.K. supervised the project. A.B. performed genome assembly and annotation. A.B., R.M., C.N., K.L and G.K contributed to other bioinformatic analyses and data deposition. C.N., K.L., B.T. and A.F. collected and prepared the samples. All authors discussed and interpreted the data. CN wrote the draft manuscript and all authors read and approved the final manuscript.

## Literature Cited

[1] Bevan, M. W., Uauy, C., Wulff, B. B., Zhou, J., Krasileva, K., & Clark, M. D. Genomic innovation for crop improvement. Nature 543(7645), 346 (2017).

[2] Chen, F. et al. Genome sequences of horticultural plants: past, present, and future. Hortic. Res. 6(1), 1–23 (2019).

[3] International Nut and Dried Fruit Council, Statistical Yearbook 2018/2109, https://www.nutfruit.org/industry/news/detail/statistical-yearbook

[4] Christenhusz, M. J., & Byng, J. W. The number of known plants species in the world and its annual increase. Phytotaxa 261(3), 201–217 (2016).

[5] Nock, C. J., Baten, A., & King, G. J. Complete chloroplast genome of *Macadamia integrifolia* confirms the position of the Gondwanan early-diverging eudicot family Proteaceae. BMC Genomics 15(S9), S13 (2014).

[6] Hardner, C. Macadamia domestication in Hawai‘i. Genet. Resour. Crop Evol. 63(8), 1411–1430 (2016).

[7] Powell, M., Accad, A., & Shapcott, A. Where they are, why they are there, and where they are going: using niche models to assess impacts of disturbance on the distribution of three endemic rare subtropical rainforest trees of *Macadamia* (Proteaceae) species. Aust. J. Bot. 62(4), 322–334 (2014).

[8] Nock, C. J. et al. Wild origins of macadamia domestication identified through intraspecific chloroplast genome sequencing. Front. Plant Sci. 10, 334 (2019).

[9] Topp, B. L., Nock, C. J., Hardner, C. M., Alam, M., & O’Connor, K. M. Macadamia (*Macadamia* spp.) Breeding. In Advances in Plant Breeding Strategies: Nut and Beverage Crops 221–251 (Springer, 2019).

[10] Chabikwa, T. G., Barbier, F. F., Tanurdzic, M., & Beveridge, C. A. De novo transcriptome assembly and annotation for gene discovery in avocado, macadamia and mango. Sci. Data 7(1), 1–7 (2020).

[11] Lim, S. L., D’Agui, H. M., Enright, N. J., & He, T. Characterization of leaf transcriptome in Banksia hookeriana. Genom. Proteom. Bioinf. 15(1), 49–56 (2017).

[12] Damerval, C. et al., 2019. Unravelling the developmental and genetic mechanisms underpinning floral architecture in Proteaceae. Front. Plant Sci. 10, 18 (2019).

[13] Akman, M., Carlson, J. E., Holsinger, K. E., & Latimer, A. M. Transcriptome sequencing reveals population differentiation in gene expression linked to functional traits and environmental gradients in the South African shrub *Protea repens*. New Phytol. 210(1), 295–309 (2016).

[14] Ostria-Gallardo, E. et al., Transcriptomic analysis suggests a key role for SQUAMOSA PROMOTER BINDING PROTEIN LIKE, NAC and YUCCA genes in the heteroblastic development of the temperate rainforest tree Gevuina avellana (Proteaceae). New Phytol. 210(2), 694–708 (2016).

[15] O’Connor, K., Hayes, B., Hardner, C., Nock, C., Baten, A., Alam, M., Henry, R. and Topp, B. Genome-wide association studies for yield component traits in a macadamia breeding population. BMC genomics, 21(1), 1–12 (2020).

[16] Nock, C.J., Baten, A., Barkla, B.J., Furtado, A., Henry, R.J. & King, G.J., Genome and transcriptome sequencing characterises the gene space of *Macadamia integrifolia* (Proteaceae). BMC Genomics, 17(1), 937 (2016).

[17] Langdon, K.S., King, G.J., Baten, A., Mauleon, R., Bundock, P.C., Topp, B.L. and Nock, C.J. Maximising recombination across macadamia populations to generate linkage maps for genome anchoring. Scientific Reports, 10(1), 1–15 (2020).

[18] Furtado, A. (2014) DNA Extraction from Vegetative Tissue for Next-Generation Sequencing. In: Cereal Genomics vol. 1099 (Henry, R. J. and Furtado, A., eds.), pp. 1–5: (Humana Press, 2014).

[19] Andrews, S. FastQC: a quality control tool for high throughput sequence data, http://www.bioinformatics.babraham.ac.uk/projects/fastqc (2010).

[20] Bushnell, B., 2014. BBMap: a fast, accurate, splice-aware aligner (No. LBNL-7065E). (Lawrence Berkeley National Lab, 2014) https://jgi.doe.gov/data-and-tools/bbtools/bb-tools-user-guide/bbmap-guide/

[21] Salmela, L., & Rivals, E. LoRDEC: accurate and efficient long read error correction. Bioinformatics 30, 3506–3514 (2014).

[22] Zimin, A. V. et al. The MaSuRCA genome assembler. Bioinformatics 29, 2669–2677 (2013).

[23] Boetzer, M., Henkel, C.V., Jansen, H.J., Butler, D. & Pirovano, W. Scaffolding pre-assembled contigs using SSPACE. Bioinformatics, 27, 578–579 (2011).

[24] Xue W. et al. L RNA scaffolder: scaffolding genomes with transcripts. BMC Genomics 2013;14(1):604.

[25] Grabherr, M. G. et al. Full-length transcriptome assembly from RNA-seq data without a reference genome. Nat. Biotechnol., 29, 644–652 (2011).

[26] Tang, H. et al. 2015. ALLMAPS: robust scaffold ordering based on multiple maps. Genome Biol., 16(1), 3 (2015).

[27] Marcais, G. & Kingsford, C. A fast, lock-free approach for efficient parallel counting of occurrences of k-mers. Bioinformatics, 27(6), 764–70 (2011).

[28] Sun, H., Ding, J., Piednoël, M. and Schneeberger, K., findGSE: estimating genome size variation within human and Arabidopsis using k-mer frequencies. Bioinformatics, 34(4), 550–557 (2018).

[29] Vurture, G. W. et al. GenomeScope: fast reference-free genome profiling from short reads. Bioinformatics, 33(14), 2202–2204 (2017).

[30] Li, H., 2018. Minimap2: pairwise alignment for nucleotide sequences. Bioinformatics, 34(18), 3094–3100 (2018).

[31] Van der Auwera GA, Carneiro M, Hartl C, Poplin R, del Angel G, Levy-Moonshine A, Jordan T, Shakir K, Roazen D, Thibault J, Banks E, Garimella K, Altshuler D, Gabriel S, DePristo M. (2013) From FastQ Data to High-Confidence Variant Calls: The Genome Analysis Toolkit Best Practices Pipeline. Current Protocols In Bioinformatics, 43:11.10.1–11.10.33 (2013).

[32] Tarailo-Graovac, M., & Chen, N. Using RepeatMasker to identify repetitive elements in genomic sequences. Curr. Protoc. Bioinformatics, 25(1), 4–10 (2009).

[33] Campbell, M. S., Holt, C., Moore, B., & Yandell, M. (2014). Genome annotation and curation using MAKER and MAKER-P. Curr. Protoc. Bioinformatics, 48(1), 4–11.

[34] Xu, L., Dong, Z., Fang, L., Luo, Y., Wei, Z., Guo, H., Zhang, G., Gu, Y.Q., Coleman-Derr, D., Xia, Q. and Wang, Y., 2019. OrthoVenn2: a web server for whole-genome comparison and annotation of orthologous clusters across multiple species. Nucleic acids research, 47(W1), pp.W52–W58.

[35] Simão, F. A., Waterhouse, R. M., Ioannidis, P., Kriventseva, E. V. & Zdobnov, E. M. BUSCO: assessing genome assembly and annotation completeness with single-copy orthologs. Bioinformatics 31, 3210–3212 (2015).

